# Somatomotor disconnection links sleep duration with socioeconomic context, screen time, cognition, and psychopathology

**DOI:** 10.1101/2024.10.29.620865

**Authors:** Cleanthis Michael, Aman Taxali, Mike Angstadt, Katherine L. McCurry, Alexander Weigard, Omid Kardan, M. Fiona Molloy, Katherine Toda-Thorne, Lily Burchell, Maria Dziubinski, Jason Choi, Melanie Vandersluis, Luke W. Hyde, Mary M. Heitzeg, Chandra Sripada

## Abstract

**Background:** Sleep is critical for healthy brain development and emotional wellbeing, especially during adolescence when sleep, behavior, and neurobiology are rapidly evolving. Theoretical reviews and empirical research have historically focused on how sleep influences mental health through its impact on *higher-order* brain systems. No studies have leveraged data-driven network neuroscience methods to uncover interpretable, brain-wide signatures of sleep duration in adolescence, their socio-environmental origins, or their consequences for cognition and mental health.

**Methods:** Here, we implement graph theory and component-based predictive modeling to examine how a multimodal index of sleep duration is associated with intrinsic brain architecture in 3,173 youth (11-12 years) from the Adolescent Brain Cognitive Development^SM^ Study.

**Results:** We demonstrate that network integration/segregation exhibit a strong, generalizable multivariate association with sleep duration. We next identify a single component of brain architecture centered on a single network as the dominant contributor of this relationship. This component is characterized by increasing disconnection of a *lower-order* system - the somatomotor network - from other systems, with shorter sleep duration. Finally, greater somatomotor disconnection is associated with lower socioeconomic resources, longer screen times, reduced cognitive/academic performance, and elevated externalizing problems.

**Conclusions:** These findings reveal a novel neural signature of shorter sleep in adolescence that is intertwined with environmental risk, cognition, and psychopathology. By robustly elucidating the key involvement of an understudied brain system in sleep, cognition, and psychopathology, this study can inform theoretical and translational research directions on sleep to promote neurobehavioral development and mental health during the adolescent transition.

## Introduction

Sleep is a basic human need and plays a critical role in cognition, emotion regulation, academic and occupational attainment, and physical and mental health (1). Experimental, longitudinal, genetically informed, and clinical trial designs demonstrate that sleep causally influences health and behavior across the lifespan (2–5). However, large-scale studies and meta-analyses indicate that most adolescents sleep less than recommended (6). Inadequate sleep during adolescence has been identified cross-culturally (7) and, given systemic inequities, is more common within disadvantaged communities (8), perpetuating population-level health disparities.

Most psychiatric disorders emerge during adolescence (9), a pattern that has recently worsened and theorized to partly emanate from changing trends in sleep (10). Accordingly, the National Sleep Foundation has declared sleep deprivation during adolescence as a “public health epidemic” (11). Numerous factors unique to adolescence may shorten sleep, such as increasing academic and social demands, longer screen times, and circadian and physiological remodeling, coupled with early school start times (12). These considerations underscore the importance of delineating the biological mechanisms through which sleep duration influences health and behavior during adolescence, which can inform early risk identification, policy, and intervention.

Cross-species evidence demonstrates that a central function of sleep is to promote healthy brain development (13). Indeed, empirical research has documented associations between adolescent sleep duration and quality with the structure and function of individual brain regions (12). The majority of these studies have adopted a theoretically-driven approach, examining an *a priori* subset of regions underlying higher-order processes such as executive functions, emotion regulation, decision-making, learning, and memory (e.g., prefrontal cortex, amygdala, striatum, hippocampus) (14). This literature has begun to illuminate how normative and extreme variations in adolescent sleep patterns become neurobiologically and behaviorally expressed.

Recently, there has been increasing recognition that the brain is a complex network, such that coordinated activity across the entire brain gives rise to cognitive and affective processes that shape risk for psychiatric illness (15,16). Technological and computational advancements in non-invasive neuroimaging methods provide novel insight into how information is processed across the brain. Specifically, task-free “resting-state” functional magnetic resonance imaging (fMRI) uses coherence in spontaneous neural activity to map the strength of communication between brain regions (functional connectivity) (15,16). Seminal resting-state work has revealed that the brain organizes into intrinsic connectivity networks (ICNs) that support diverse functions (17). Accordingly, examining large-scale ICNs, rather than only individual regions, may be central to understanding how sleep duration calibrates adolescent neurobehavioral development.

Advanced techniques from network neuroscience, such as graph theory, offer computational tools to quantify parsimonious and interpretable properties of how ICNs are organized across the brain (18). Across adolescence, ICNs exhibit complex developmental sequences in segregation (communication within distinct ICNs) and integration (communication across different ICNs) (19–21). Segregation yields differentiated networks that execute specialized functions, while integration efficiently coordinates these processing streams across the brain (22). Segregation and integration are reflected in two graph theoretic metrics that capture within-network (within-module degree) and between-network connectivity (participation coefficient) (23). Thus, graph properties of segregation/integration can provide a parsimonious and developmentally informed account of how sleep influences brain and behavior during adolescence.

Several lines of evidence suggest that normative patterns of segregation/integration may be perturbed by shortened sleep in adolescence. Sleep is intertwined with multiple environmental stressors (or resources) that have been associated with ICN organization, such as household instability (24) and socioeconomic resources (25–27). Further, sleep patterns can impact developmental processes that partly emerge from segregation and integration, such as psychiatric disorders, executive functioning, and multi-domain resilience (28–30). Lastly, initial studies are beginning to identify links between sleep patterns and the segregation/integration of large-scale networks in adolescence (31,32). However, no study has integrated a large sample, multi-method sleep assessments, and graph analyses to identify robust neural signatures of sleep duration in adolescence. Additionally, how these neural signatures relate to associated risk factors (e.g., socioeconomic context) and translate into cognition and psychopathology is unknown.

The present study therefore sought to identify robust graph theoretic signatures of shorter sleep, and their relationship with context and behavior. We leveraged the Adolescent Brain Cognitive Development^SM^ (ABCD®) Study, a large population-based consortium study of early adolescents with substantial sociodemographic diversity (33). Given disagreements between self-reported and actigraphic measures of sleep duration (34), we triangulated objective and subjective sleep assessments across multiple reporters (child-report, parent-report, Fitbit) to precisely characterize sleep duration and its link with intrinsic network segregation/integration.

Using multivariate predictive modeling, we establish that sleep is robustly related to segregation and integration across the entire brain. Contrary to prior research focusing on how sleep impacts higher-order ICNs (12), we demonstrate that disconnection of the *somatomotor* network represents a robust neural signature of shorter sleep. This somatomotor-centered motif is also linked to key developmental contexts (socioeconomic resources, screen use) and outcomes (psychopathology, cognition). Our study highlights the utility of data-driven, brain-wide approaches for elucidating novel neural signatures, and potential intervention targets, of shorter sleep during adolescence.

## Methods and Materials

### Participants

The ABCD Study is a longitudinal study with 11,875 children (9-10 years) from 22 sites across the US. The study conforms to procedures of each site’s Institutional Review Board. Participants provide informed consent (parents) or assent (children). This study uses ABCD Release 4.0 data.

### Functional Neuroimaging

High spatial (2.4mm isotropic) and temporal resolution (TR=800ms) resting-state fMRI was acquired in four 5-minute runs (20min total). Preprocessing was performed using fMRIPrep v1.5.0 (35) and included surface reconstruction (FreeSurfer v6.0.01), spatial normalization, rigid co-registration to the T1-weighted image, motion correction, normalization to standard space, and transformation to CIFTI space.

Connectomes were generated for each resting-state run using the Gordon-333 cortical atlas (17), augmented with high-resolution subcortical (36) and cerebellar (37) atlases. Volumes exceeding a framewise displacement (FD) threshold of 0.5mm were censored. Covariates were regressed out of the timeseries in a single step, including: linear trend, 24 motion parameters (original translations/rotations, derivatives, quadratics), aCompCorr 5 CSF and 5 WM components and aggressive ICA-AROMA components, high-pass filtering (0.008Hz), and censored volumes. Next, correlation matrices were calculated (**Supplement** and **fMRIPrep Supplement**).

### Inclusion/Exclusion

ABCD Release 4.0 includes 11,875 participants, with neuroimaging data at baseline (ages 9-10) and year-2 (ages 11-12). Our primary analysis focused on year-2 data, which has three measures of sleep duration. Participants were excluded for: failing ABCD quality control (QC), insufficient number of resting-state runs each ≥4 minutes after censoring frames with FD>0.5mm, failing visual QC of registrations and normalizations, and missing data required for analysis (**Supplement**). This left *N*=5,596 participants across 21 sites for our main principal component analysis identifying components of brain architecture (“somatomotor disconnection”; **Results**), and *N*=3,173 participants with complete data on all sleep measures for examining sleep-brain associations. See **Supplement** for sample sizes for analyses evaluating brain-context and brain-phenotype effects (**Socio-Environmental Context and Behavioral Phenotypes**).

### Graph Theory

As most graph measures require unsigned edge weights, we focused on positive connections consistent with previous graph theoretical investigations (38,39).

#### Network Segregation

We calculated *within-module degree*, a node-wise measure that captures each node’s connectivity strength within its own network. This metric modifies the “module-degree z-score” metric (23) by bypassing within-network z-scoring to better capture differences across participants, rather than differences across nodes within each network.

#### Network Integration

We calculated *participation coefficient*, a node-wise measure that captures the diversity of a node’s connections with nodes outside its own network (23). If a node distributes its connectivity evenly across networks, its participation coefficient will be 1, while equality departures yield commensurately lower scores.

We used the community structure defined by the applied parcellation schemes to determine network boundaries. Within-module degree (MDP) and participation coefficient (PCP) for positive edges were calculated for 418 nodes, yielding 836 node-wise graph theoretic features per participant.

### Sleep Duration

We constructed a latent variable for sleep duration using factor analysis of three measures (**Supplement**):

#### Sleep Disturbance Scale (SDSC)

The parent-reported SDSC includes 26 questions assessing six domains of sleep disturbances over the past six months using a 5-point Likert scale (40). This scale has been used in other studies in ABCD, validly and reliably measures sleep disturbances, and fulfills psychometric requirements among childhood sleep scales. We focused on this scale’s sleep duration item.

#### ABCD Youth Munich Chronotype Questionnaire (MCTQ-C)

The self-reported MCTQ-C includes 17 items that assess diurnal preferences that manifest in personal sleep-wake rhythms (chronotypes), including sleep/wake schedules on school and school-free days (41). This scale has been used in other studies in ABCD. We focused on the sleep duration item for school days.

#### FitBit

Fitbit devices assess biobehavioral features (sleep, physical activity) objectively, continuously, and unobtrusively. Youth wore a Charge 2 Fitbit over 21 days and were instructed to remove it only when charging and bathing (42). The Fitbit app was downloaded on the youth or parent’s phone, and participants were instructed to sync the Fitbit daily, monitoring data using *Fitabase*. Sleep intervals were captured by an intrinsic device algorithm. Fitbit measures of sleep have been examined in other studies in ABCD and demonstrate adequate sensitivity (43).

### Multivariate Predictive Modeling

To quantify the multivariate relationship between these 836 graph metrics and sleep duration, we used principal component regression (PCR) predictive modeling (44,45) (**Supplement**). This method applies principal component analysis (PCA) on the predictive features (MDP/PCP), fits a regression model on the resulting components (determined in nested cross-validation), and implements this model out-of-sample using leave-one-site-out cross-validation (LOSO-CV). We control for sex assigned at birth, parent-reported race/ethnicity, age, age-squared, mean FD, and mean FD-squared. We controlled for race/ethnicity, a social construct, to account for differences in exposure to personal and systemic racism, disadvantage, and opportunity among people of color in the US (46). Significance was determined with non-parametric permutation tests, using the procedure of Freedman and Lane (47) to account for covariates. Exchangeability blocks accounted for twin, family, and site structure by entering them into Permutation Analysis of Linear Models (48) to produce permutation orderings. We visualize the multivariate model using a standardized importance map, which multiplies derived principal components by their standardized beta weights, followed by z-scoring to facilitate interpretation.

### Socio-Environmental Context and Behavioral Phenotypes

Lastly, we examined the potential origins and consequences of somatomotor disconnection (**Supplement**). Briefly, we investigated key developmental contexts including neighborhood disadvantage (area deprivation index), household income, parental education, and screen time, and key developmental outcomes including youth-reported (Brief Problem Monitor) and parent-reported (Child Behavior Checklist) externalizing and internalizing symptoms, general cognitive ability (latent variable from ABCD neurocognitive battery), and grades (School Environment subscale of the School Risk and Protective Factors Survey). We conducted linear regression models predicting each contextual/phenotypic variable from somatomotor disconnection, controlling for sex, race/ethnicity, age, age-squared, mean FD, mean FD-squared, and site.

## Results

### Robust associations between shorter sleep duration and network segregation/integration

We initially predicted shorter sleep duration from 836 indicators of node-wise network segregation/integration (MDP/PCP) in the year-2 data using LOSO-CV to quantify predictivity and ensure generalizability to new participants. This out-of-sample multivariate relationship was r_cv_= 0.220, p_PERM_ < 0.0001. **Figure 1** depicts a standardized importance map, which captures the graph theoretic predictors of sleep duration, weighted by their importance. This map illustrates a prominent role for both the segregation and integration of somatomotor nodes, which exhibited higher segregation (MDP) and lower integration (PCP) with shorter sleep duration. We describe this architectural motif as “somatomotor disconnection”.

**Figure 1.**
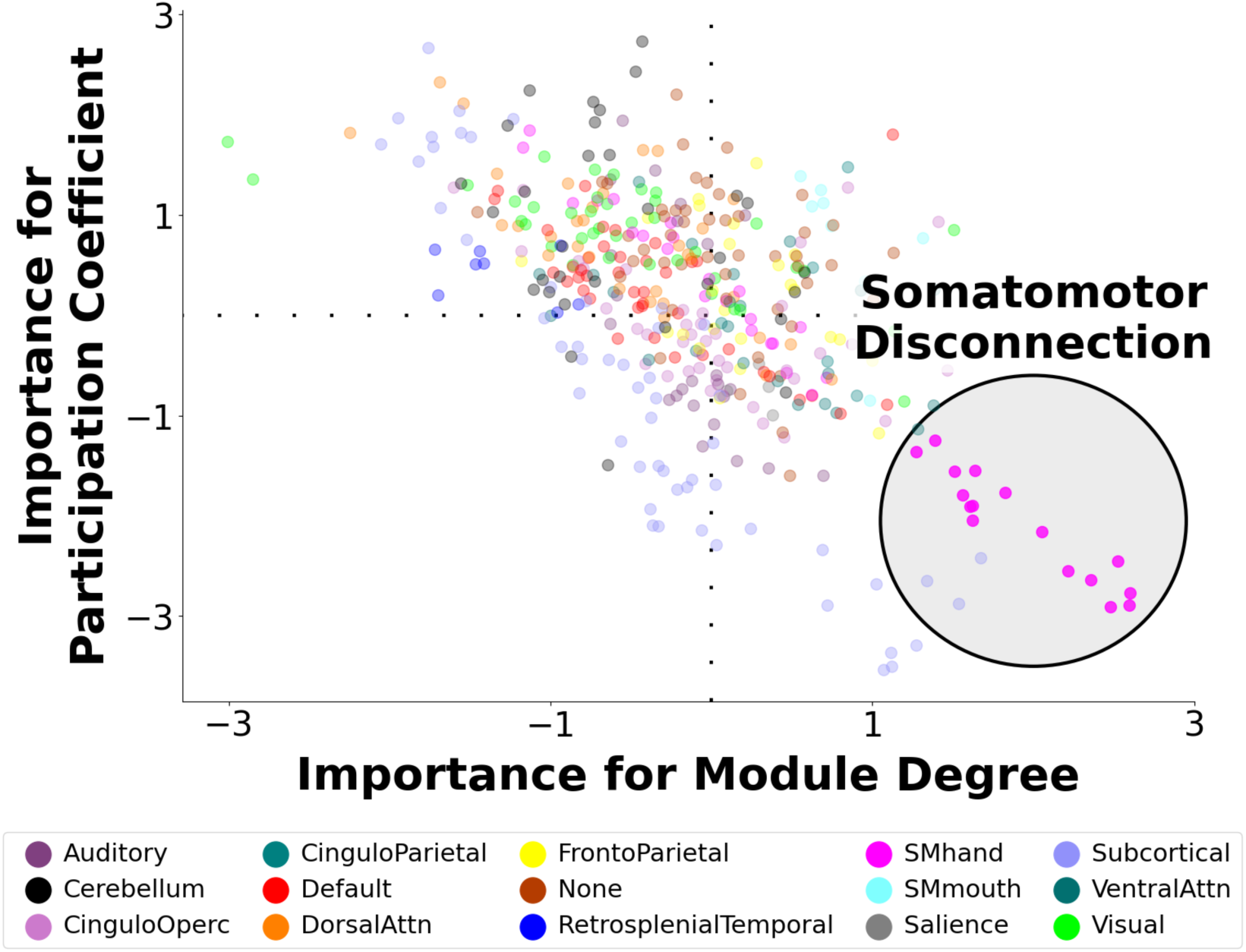
Multivariate Neural Signature of Shorter Sleep Duration. A multivariate predictive model was trained to predict shorter sleep duration in youth using 836 graph theoretic metrics capturing network segregation (within-module degree) and network integration (participation coefficient) during rest. In leave-one-site-out cross-validation, the model robustly predicted shorter sleep duration in held-out participants. The figure depicts a visualization of the neural signature that was the basis of prediction in which each node of the brain is assigned an importance score for within-module degree for positive edges (MDP) and participation coefficient for positive edges (PCP), based on how important the respective metric was for predicting sleep duration. The sign indicates whether each node’s segregation/integration is positively or negatively related to shorter sleep duration, and the absolute value indicates the importance of the node on that dimension. This map illustrates a prominent motif of somatomotor disconnection in the lower right, in which among youth with shorter sleep duration, nodes of the somatomotor network exhibit increased within-module degree and reduced participation coefficient. This topological pattern reflects greater segregation (“disconnection”) of the somatomotor network with shorter sleep duration.

### Network segregation/integration capture almost the entire multivariate relationship between sleep duration and the full connectome

We implemented the same multivariate model to examine the relationship between shorter sleep duration and the full connectome (87,153 connections). The out-of-sample multivariate relationship of sleep duration with the entire connectome (r_cv_= 0.219, p_PERM_ < 0.0001) was slightly lower than with the 836 graph metrics.

We therefore tested whether the 836 graph metrics reflect distinct versus overlapping variance in predicting shorter sleep duration relative to the full connectome. We built a stacked model by simultaneously entering both the predictions from the MDP/PCP model, and those from the full connectome model. The out-of-sample performance of this stacked model was r_cv_= 0.231; that is, the stacked model performed similarly to the 836-feature graph theoretic model alone. These results suggest predictivity of sleep duration is strongly concentrated in the graph theoretic features, wherein these 836 MDP/PCP metrics account for most of the multivariate link between the full connectome and sleep duration, despite the 100-fold reduction in the number of features.

### A single component, which exhibits the somatomotor disconnection motif, is the dominant contributor to the sleep-brain relationship

Our predictive model approach applied PCA to 836 graph metrics in our year-2 participants, yielding components of inter-individual variation that strongly predicted sleep duration. We next examined each component’s predictivity of sleep duration with a scree plot (**Figure 2**, left panel). We found that component #3 was the dominant contributor in predicting sleep duration. This component demonstrated a prominent somatomotor disconnection motif (**Figure 2**, right panel). The standardized regression coefficient of this somatomotor disconnection component (controlling for sex, race/ethnicity, age, age-squared, mean FD, mean FD-squared, and site) with sleep duration is 0.15, 68% as strong as the main model using 59 graph theoretic components.

**Figure 2.**
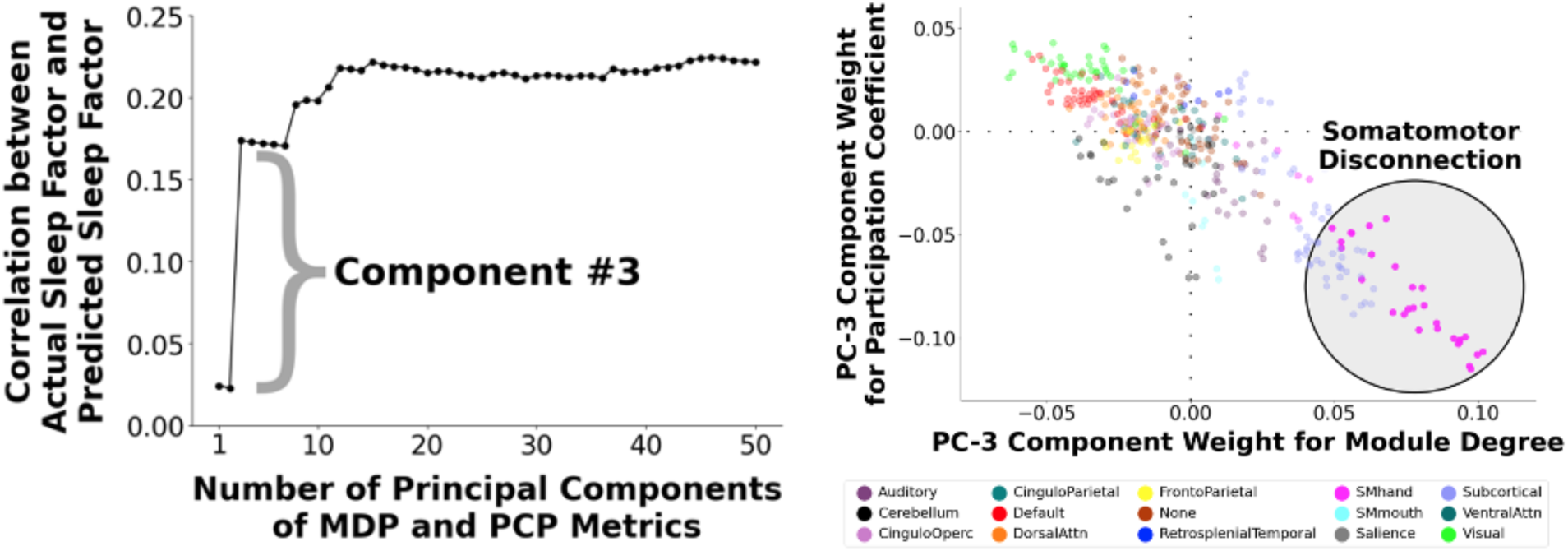
(Left) Contributions of Individual Graph Theory Principal Components (PC) to the Multivariate Prediction of Shorter Sleep Duration. We conducted a multivariate predictive model to predict shorter sleep duration from 836 node-wise metrics reflecting network segregation (within-module degree for positive connections; MDP) and network integration (participation coefficient for positive connections; PCP). We constructed a scree plot to examine contributions of individual components to the multivariate prediction of shorter sleep duration. Component #3 was found to be the dominant contributor in predicting shorter sleep duration. **(Right) Visualization of Component #3.** We visualized the feature weights for component #3. This component displayed a prominent somatomotor disconnection motif in which the nodes of the somatomotor network exhibit higher within-module degree and lower participation coefficient (i.e., higher segregation, or “disconnection”, relative to other brain networks).

We previously showed that PCA-derived brain components serve as “basic units” of neural variation that stably and systematically differ across individuals (44). We therefore assessed the robustness of our results by examining how shorter sleep duration relates to component #3 derived from an independent sample. From the ABCD baseline sample with good resting-state data (*N*=6,570), we identified 3,148 participants who did not overlap with those in our main analysis that used year-2 data, and repeated our models, again focusing on the somatomotor disconnection component. Despite being derived from a non-overlapping sample, the baseline- and year-2-derived components #3 were nearly identical, exhibiting a prominent somatomotor disconnection motif (**Supplement**) and highly correlated (r=0.99) feature weights.

Second, we calculated the expression of this baseline-derived component #3 for each participant in our year-2 sample by projecting each year-2 participant’s graph metrics onto the coefficients for this component via a vector dot product. The standardized regression coefficient of these scores (controlling for sex, race/ethnicity, age, age-squared, mean FD, mean FD-squared, and site) with sleep duration was 0.17, similar to the relationship with the year 2-derived component, supporting the robustness of this component.

### Sleep-related somatomotor disconnection is associated with key developmental contexts and outcomes

We next examined how somatomotor disconnection was related to socioeconomic context and behavioral phenotypes, controlling for sex, race/ethnicity, age, age-squared, mean FD, mean FD-squared, and site. We focused on the baseline-derived somatomotor disconnection component, ensuring the samples used to generate somatomotor disconnection versus to evaluate its environmental-phenotypic associations are independent. Since baseline- and year 2-derived components were correlated at 0.99, results using the year-2-derived component were similar.

As demonstrated in **Figure 3**, somatomotor disconnection was related to key developmental contexts, including household income (β=-0.05, *p*=1.0×10^−4^), parental education (β=-0.09, *p*=5.7×10^−11^), area deprivation index (β=0.03, *p*=0.003), and screen time (β=0.13, *p*=1.5×10^−19^). Somatomotor disconnection was also associated with key developmental outcomes, including youth-reported (β=0.06, *p*=4.5×10^−4^) and parent-reported externalizing symptoms (β=0.05, *p*=0.001), school grades (β=-0.04, *p*=0.007), and general cognitive ability (β=-0.06, *p*=4.3×10^−5^), but not youth-reported (β=0.03, *p*=0.103) or parent-reported (β=-0.01, *p*=0.472) internalizing symptoms.

**Figure 3.**
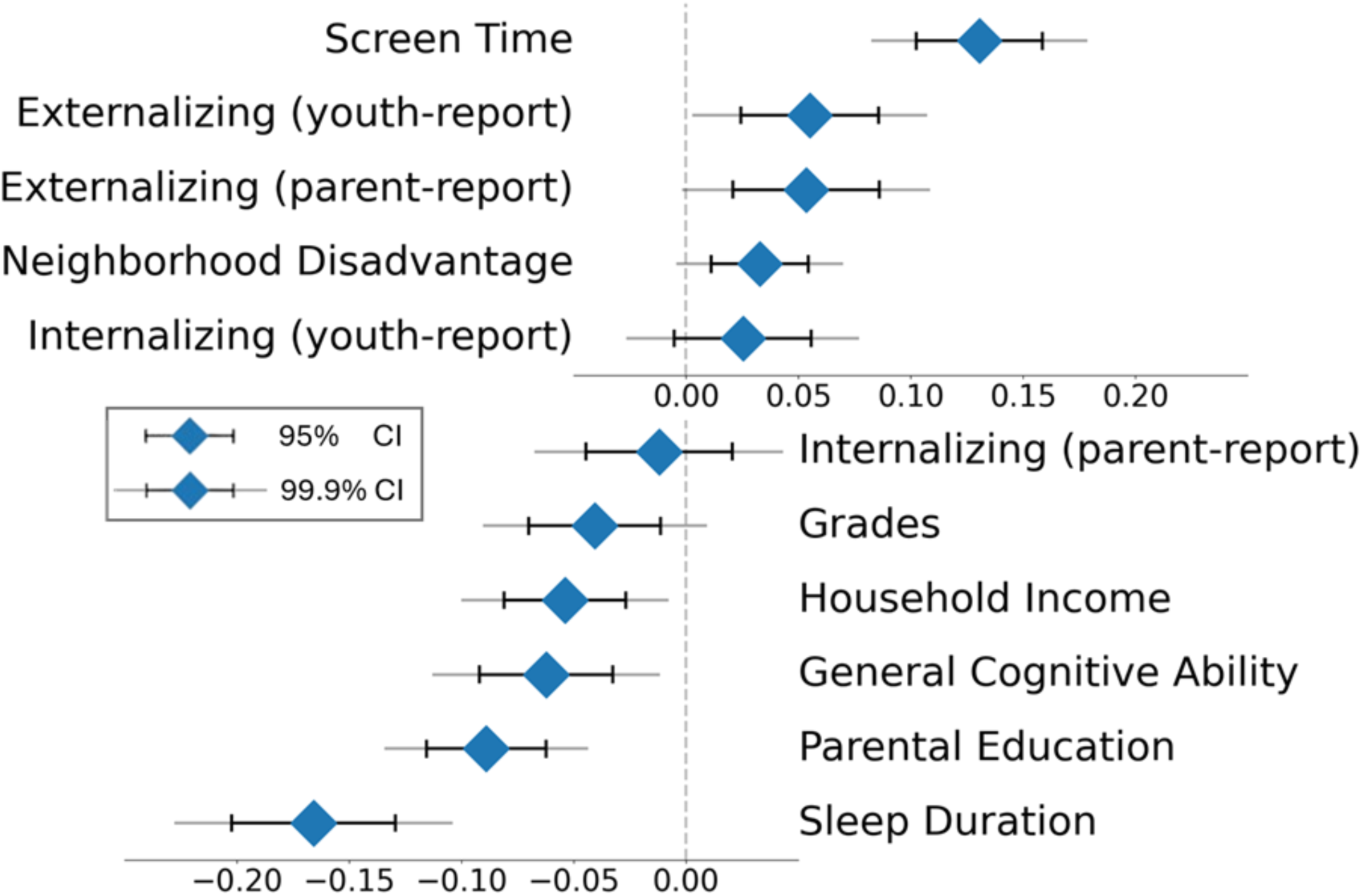
Associations of the Sleep-Related Somatomotor Disconnection Component with Socio-Environmental and Behavioral Phenotypes. Greater “disconnection” of the somatomotor network, which was strongly related to shorter sleep duration, was significantly associated with longer screen times, higher youth-reported externalizing problems (Brief Problem Monitor Form), higher parent-reported externalizing problems (Child Behavior Checklist), higher neighborhood disadvantage (area deprivation index), lower school grades, lower household income, general cognitive ability (latent variable from ABCD neurocognitive battery scores), and lower parental education. Sleep-related somatomotor disconnection was not significantly associated with youth-(Brief Problem Monitor Form) or parent-reported (Child Behavior Checklist) internalizing problems. Diamonds represent the mean, black lines represent 95% confidence intervals, and gray lines represent 99.9% confidence intervals.

Finally, we tested the association between somatomotor disconnection and shorter sleep duration when controlling for the preceding socio-environmental and behavioral variables. The relation remained significant (β=-0.13, *p*=4.9×10^−9^), indicating that somatomotor disconnection shares variance with socioeconomic context, behavior, and cognition, but is uniquely linked to sleep duration, even after accounting for intertwined exposures and phenotypes.

## Discussion

Shorter sleep increases risk for impaired regulation over behavior, cognition, and emotion (1,3,5), is especially common in adolescence (6), disrupts brain development (12), and incurs substantial societal and financial costs (49). No study has identified parsimonious brain signatures underlying shortened sleep and its neurobehavioral sequelae during this critical developmental period. By integrating techniques from network science, latent variable modeling across brain and behavior, and multivariate predictive analyses in a large, sociodemographically diverse cohort, we elucidate a novel neural signature of shorter sleep in adolescence. Specifically, we demonstrate a robust and generalizable link between reduced sleep and disconnection of the somatomotor network at rest. This sleep-related architectural motif was further associated with socioeconomic context, screen time, cognition, and psychopathology. This brain signature could serve as a novel neural marker for early risk identification, extends our understanding of how environmental insults confer risk, and may inform the development of new psychosocial and neuromodulatory interventions.

### Multivariate link between sleep duration and network segregation/integration

Given disagreements between subjective and objective measures of sleep (34), we triangulated subjective (self- and parent-report) and objective (Fitbit) assessments to construct a multimodal index of sleep duration. Similar to prior work (50), the out-of-sample correlation between sleep duration and the full connectome was 0.22. While this effect is relatively large for context-brain associations, multivariate techniques *describing* how sleep duration relates to 87,153 connections do not afford clear interpretations of *what* these neural alterations mean and *how* they are organized across the brain.

Here, we leveraged graph theory to distill these 87,153 connections into only 836 features with neuroscientific interpretability. We quantified each region’s integration (participation coefficient) and segregation (within-module degree). Despite this 100-fold reduction in number of features, we still captured almost the entire association between sleep duration and brain organization. These findings suggest that network segregation/integration offer a parsimonious and interpretable framework of how adolescent sleep duration relates to neural architecture.

### Somatomotor disconnection as a neural signature of shorter sleep

The current literature has predominantly focused on how sleep impacts the development of *higher-order* brain systems. These systems include the default mode network, which underlies social and self-related processing (31,51,52), and its interactions with networks underlying cognitive control (fronto-parietal, dorsal attention) (52). Other studies implicate cortico-limbic circuitry that supports emotional learning and regulation (53), suggesting that sleep behaviors may primarily modulate higher-order association systems.

Against this backdrop, we expected that sleep duration would mainly calibrate the topology of association systems. Surprisingly, out of 59 components of brain architecture, one component was the dominant contributor to the multivariate association between sleep duration and brain-wide organization. The defining characteristics of this component involved high segregation (within-module degree) and low integration (participation coefficient) of a *lower-order* system: the somatomotor network, spanning precentral/postcentral gyri extending into supplementary motor cortex. We describe this motif as *somatomotor disconnection*. That is, with shorter sleep, regions in the somatomotor network became more interconnected, and less connected with other systems, making them more “disconnected” in the whole-brain architecture. We replicated this result using a somatomotor disconnection component derived from an independent set of participants and while controlling for related contexts (socioeconomic resources) and phenotypes (cognition, psychopathology), confirming the internal structure, developmental stability, and generalizability of somatomotor disconnection.

Converging evidence from innovative techniques such as graph theory (25,26,29), poly-neuro risk profiles (54), and multivariate predictive modeling (45,55) demonstrates that context and phenotypes are associated with the organization of the entire brain. Therefore, it is highly uncommon, and thus particularly remarkable, that a single component centered on a *single network* captures the bulk of individual differences in sleep duration. Notably, this component primarily involved a network not commonly considered in the literature. Accordingly, while theoretically-driven approaches delineate the role of specific brain systems, our findings illustrate that data-driven brain-wide techniques can uncover the central role of systems that have previously been understudied.

The central involvement of the somatomotor network in sleep may initially seem surprising. However, inhibitory neurotransmission in the substantia nigra, a subcortical structure critical for motor control, regulates sleep in rodents (56). Sleep quality has also been associated with intrinsic connectivity of precentral and supplementary motor areas (50,57). Lastly, sleep quality and duration have been linked with the integration of the somatomotor network (32). This literature reinforces our findings indicating primary associations between sleep duration and somatomotor architecture.

We offer several speculations about the neurophysiological implications of somatomotor disconnection. First, key motor areas such as the basal ganglia regulate sleep via neurochemical and electrophysiological mechanisms (58). Thus, somatomotor dysconnectivity may indicate impaired regulation over motor and sleep activity. Second, sleep and wakeful states differ in levels of motor activity, which may become expressed specifically along the somatomotor network. Third, neuroplasticity is highest in sensory-motor systems earlier in life and association systems later in adolescence (59). Accordingly, sleep, like other experiences, may most strongly relate to somatomotor systems earlier in development, but association systems later on (60). Additional research is required to parse whether somatomotor disconnection constitutes a cause, marker, risk factor, or consequence of shorter sleep during adolescence.

### Contextual and phenotypic implications of somatomotor disconnection

We additionally characterized how sleep-related somatomotor disconnection relates to critical developmental contexts and outcomes. We found stronger somatomotor disconnection in youth who spent more time on their screens. One interpretation is that longer screen times sacrifice time from sleep (12), magnifying sleep-related neural signatures. Moreover, we observed stronger somatomotor disconnection in youth with higher neighborhood disadvantage, lower household income, and lower parental education. This finding accords with evidence of poorer sleep in disadvantaged communities, partly due to higher average exposure to stress or ambient noise (61), and with consistent links among somatomotor connectivity and multiple forms of stress exposure (25,27,62). For example, we recently demonstrated that lower socioeconomic resources are associated with greater somatomotor segregation in the ABCD Study (27). These findings suggest that somatomotor disconnection may represent a robust neural marker of diverse forms of environmental risk and opportunity.

Greater sleep-related somatomotor disconnection was further linked to lower general cognition and school grades. While cognition is generally attributed to higher-order association systems, cognitive functions are emergent properties of activity across the entire brain, including lower-order systems (19,63). Across development, the somatomotor network becomes more integrated with other systems to coordinate specialized neural computations required for complex cognition (20,21). Precision mapping techniques also indicate that the somatomotor network interacts with the cingulo-opercular network to translate abstract cognitive representations into goal-relevant behavior (64). Finally, somatomotor dysconnectivity has been related to cognitive dysfunction in adults (65). Thus, our results suggest that somatomotor disconnection linked to shorter sleep may undermine cognitive performance in experimental and naturalistic settings.

Lastly, dovetailing with evidence that disrupted sleep heightens psychiatric vulnerability (1,3,4), we found that greater somatomotor disconnection was related to elevated externalizing symptoms. Individual and meta-analytic studies have linked somatomotor function to not only externalizing, but also internalizing and general psychopathology across the lifespan (66,67). Somatomotor dysconnectivity has been theorized to reflect motor impulsiveness that confers shared liability for diverse forms of externalizing behavior (66). Motor systems have also been implicated in several transdiagnostic processes, such as emotion regulation (68) and impulsivity (65). Additional research is required to uncover the mechanisms through which sleep-related somatomotor disconnection relates to cognition and psychopathology, and how it can be leveraged by intervention to promote wellbeing.

### Limitations and Future Directions

Limitations and recommendations for future research are important to consider. First, we focused on sleep duration given its clear implications for policy and prevention. However, other aspects of sleep (e.g., quality, consistency) are important to examine in future studies. Second, because two sleep duration measures were only available in year-2 data, our main analyses focused on that timepoint and were thus cross-sectional, restricting inferences about direction of causality or patterns of neurodevelopment. Future longitudinal investigations across longer timescales should evaluate whether somatomotor disconnection precedes or follows shorter sleep, and how it unfolds and translates into behavior across development. Third, the extent to which our results generalize across community populations with different sociodemographic backgrounds, and clinical populations with more severe sleep disruptions, remains unclear (69,70). Lastly, the strength of associations between somatomotor disconnection with phenotypic outcomes was relatively modest, highlighting that other topological patterns are also important for health and behavior.

## Conclusions

The present study leverages latent variable modeling across contextual, neurobiological, and phenotypic levels in a large neuroimaging sample to uncover a novel, generalizable, and robust neural signature of shortened sleep during adolescence. Specifically, shorter sleep duration was related to greater expression of a topological motif primarily defined by disconnection of the somatomotor network, which was in turn linked to lower socioeconomic resources, longer screen times, reduced cognitive and academic performance, and more externalizing problems. This study identifies neural markers of shorter sleep that can inform early risk identification and targeted interventions to scaffold healthy cognitive, emotional, and brain development in adolescence.

## Supporting information

Supplemental Methods & Results

## Acknowledgements

The ABCD Study is a multisite, longitudinal study designed to recruit more than 10,000 children aged 9-10 years and follow them over 10 years into early adulthood. A full list of supporters is available at: https://abcdstudy.org/about/federal-partners/. A listing of participating sites and a complete listing of the study investigators can be found at: https://abcdstudy.org/principal-investigators/. ABCD consortium investigators designed and implemented the study and/or provided data but did not necessarily participate in analysis or writing of this report. This manuscript reflects the views of the authors and may not reflect the opinions or views of the NIH or ABCD consortium investigators. The ABCD data repository grows and changes over time.

## CRediT Author Statement

**Cleanthis Michael:** Conceptualization, Methodology, Formal analysis, Writing - original draft, Visualization. **Aman Taxali:** Conceptualization, Methodology, Software, Validation, Formal analysis, Data curation, Visualization. **Mike Angstadt:** Conceptualization, Methodology, Software, Validation, Formal analysis, Data curation, Visualization. **Katherine L. McCurry:** Writing - review & editing. **Alexander Weigard:** Writing - review & editing. **Omid Kardan:** Writing - review & editing. **M. Fiona Molloy:** Writing - review & editing. **Katherine Toda-Thorne:** Writing - review & editing. **Lily Burchell:** Writing - review & editing. **Maria Dziubinski:** Writing - review & editing. **Jason Choi:** Writing - review & editing. **Melanie Vandersluis:** Writing - review & editing. **Luke W. Hyde:** Writing - review & editing, Project administration, Funding acquisition. **Mary M. Heitzeg:** Writing - review & editing, Project administration, Funding acquisition. **Chandra Sripada:** Conceptualization, Methodology, Writing - original draft, Visualization, Project administration, Funding acquisition.

## Data Availability

The present project used data from the Adolescent Brain Cognitive Development (ABCD) Study, an open-source dataset of >10,000 youth followed longitudinally to understand brain and behavioral development across adolescence. The data used in this report came from the National Institute of Mental Health Data Archive (NDA) Study 1299, 10.15154/1523041 (https://nda.nih.gov/study.html?id=1299), and the data used in our analyses can be found at NDA DOI: 10.15154/dhr7-5w50. The code for the analyses presented is publicly available through a Github repository (https://github.com/SripadaLab/ABCD_Resting_Sleep_GraphTheory).

