## Supplemental Methods & Results for "Somatomotor disconnection links sleep duration with socioeconomic context, screen time, cognition, and psychopathology"

### Supplemental Methods and Results

#### 1. Sample and Data

The ABCD study is a multisite longitudinal study with 11,875 children between 9-10 years of age from 22 sites across the United States. The study conforms to the rules and procedures of each site's Institutional Review Board, and all participants provide informed consent (parents) or assent (children). Data for this study are from ABCD Release 4.0.

#### 2. Data Acquisition, fMRI Preprocessing, and Connectome Generation

Imaging protocols were harmonized across sites and scanners. High spatial (2.4mm isotropic) and temporal resolution (TR = 800ms) resting-state fMRI was acquired in four separate runs (5min per run, 20min total). The entire data pipeline described below was run through automated scripts on the University of Michigan's high-performance cluster.

Preprocessing was performed using fMRIPrep version 1.5.0 (1), and detailed methods automatically generated by fMRIPrep software are provided in the fMRIPrep Supplement. T1-weighted (T1w) and T2-weighted images were run through recon-all using FreeSurfer v6.0.1. T1w images were also spatially normalized nonlinearly to MNI152NLin6Asym space using ANTs 2.2.0. Each functional run was corrected for fieldmap distortions, rigidly coregistered to the T1, motion corrected, and normalized to standard space. ICA-AROMA was run to generate aggressive noise regressors. Anatomical CompCor was run and the top 5 principal components of both CSF and white matter were retained. Functional data were transformed to CIFTI space using HCP's Connectome Workbench. All preprocessed data were visually inspected at two separate stages to ensure only high-quality data was included: after co-registration of the functional data to the structural data and after registration of the functional data to MNI template space.

Connectomes were generated for each functional run using the Gordon 333 parcel atlas (2), augmented with parcels from high-resolution subcortical (3) and cerebellar (4) atlases. Volumes exceeding a framewise displacement threshold of 0.5mm were marked to be censored. Covariates were regressed out of the time series in a single step, including: linear trend, 24 motion parameters (original translations/rotations + derivatives + quadratics), aCompCorr 5 CSF and 5 WM components and ICA-AROMA aggressive components, high-pass filtering at 0.008Hz, and censored volumes. Next, correlation matrices were calculated for each run. Each matrix was then Fisher r-to-z transformed, and then averaged across runs for each participant to yield their final connectome.

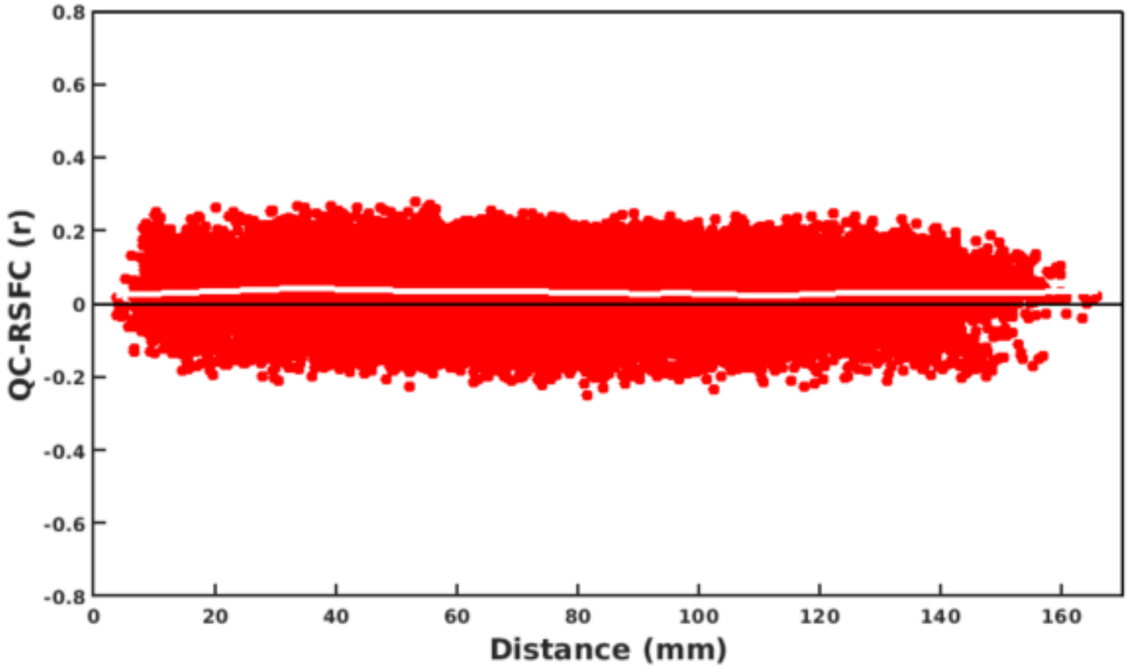


***Figure S1. Quality Control Resting-State Functional Connectivity Plot***

We used multiple procedures listed above to limit the effect of head motion on resting-state functional connectivity maps. To assess the effectiveness of these procedures, we produced a quality control resting-state functional connectivity (QC-RSFC) plot (5,6). This plot shows the relationship between mean framewise displacement and connectivity edges binned by distance. Motion effects produce a sloped line (distance-dependent artifact), while a flat line is indicative of minimal motion-related effects. The QC-RSFC plot for our ABCD resting-state data showed a flat line (Figure S1), providing additional evidence that our stringent motion correction strategies were effective.

#### 3. Inclusion/Exclusion

There are 11,875 participants in the ABCD Release 4.0 dataset. Screening was initially done using ABCD raw QC to limit to participants with 2 or more good runs of resting data as well as a good T1 and T2 image (QC score, protocol compliance score, and complete all = 1). Each run was visually inspected for registration and warping quality, and only those participants who still had 2 or more good runs were retained. After connectome generation, runs were excluded if they had less than 4 minutes of uncensored data, and next participants were retained only if they had 2 or more good runs (N = 5,596). Finally, participants who were missing data required for factor modeling of sleep were dropped from that analysis, leaving N = 3,173. Demographic characteristics of the excluded and included sample are shown in Table S1.

| | Included Sample |
|---|---|
| N | 3173 |
| Age (mean (s.d.)) | 11.95 (0.65) |
| Female (%) | 1596 (50.3) |
| Race-Ethnicity (%) | |
| non-Hispanic White | 1905 (60.0) |
| non-Hispanic Black | 276 (8.7) |
| Hispanic | 622 (19.6) |
| non-Hispanic Asian | 58 (1.8) |
| Multi-racial/Other | 312 (9.8) |
| No answer | – |
| Average Parental Education (%) | |
| < HS Diploma | 81 (2.6) |
| HS Diploma/GED | 218 (6.9) |
| Some College | 752 (23.7) |
| Bachelor | 889 (28.0) |
| Post Graduate Degree | 1229 (38.7) |
| No answer | 4 (0.1) |
| Household Marital Status – Married (%) | 2310 (72.8) |
| Household Income (%) | |
| <50K | 646 (20.4) |
| >=50k & <100K | 842 (26.5) |
| >=100k | 1570 (49.5) |
| No answer | 115 (3.6) |

***Table S1. Demographic Characteristics of Included Versus Excluded Participants***

#### 4. Graph Theoretic Measures of Functional Brain Architecture

For each participant, we calculated network segregation (within-module degree) and network integration (participation coefficient) from their weighted functional connectivity matrices, focusing on positive connections.

Within-module degree is a graph theoretic measure of within-network connectivity. Formally, the within-module degree of a node *i* is given by:

$$\sum_{j=1}^{N_i} e_{ij}$$

where $e_{ij}$ is the edge weight between nodes $i$ and $j$, and $N_i$ is the set of nodes incident to node $i$ that are in the same network as *i*.

Participation coefficient is a graph theoretic measure of between-network connectivity. Formally, the participation coefficient of a node *i* is given by:

$$1 - \sum_{m}^{M} \left(\frac{e_i(m)}{e_i}\right)^2$$

where $M$ is the set of networks, $e_i(m)$ is the sum of edge weights between node $i$ and all nodes in network $m$ and $e_i$ is the sum of edge weights between node $i$ and all other nodes.

#### 5. Principal Components Regression-Based Multivariate Predictive Modeling

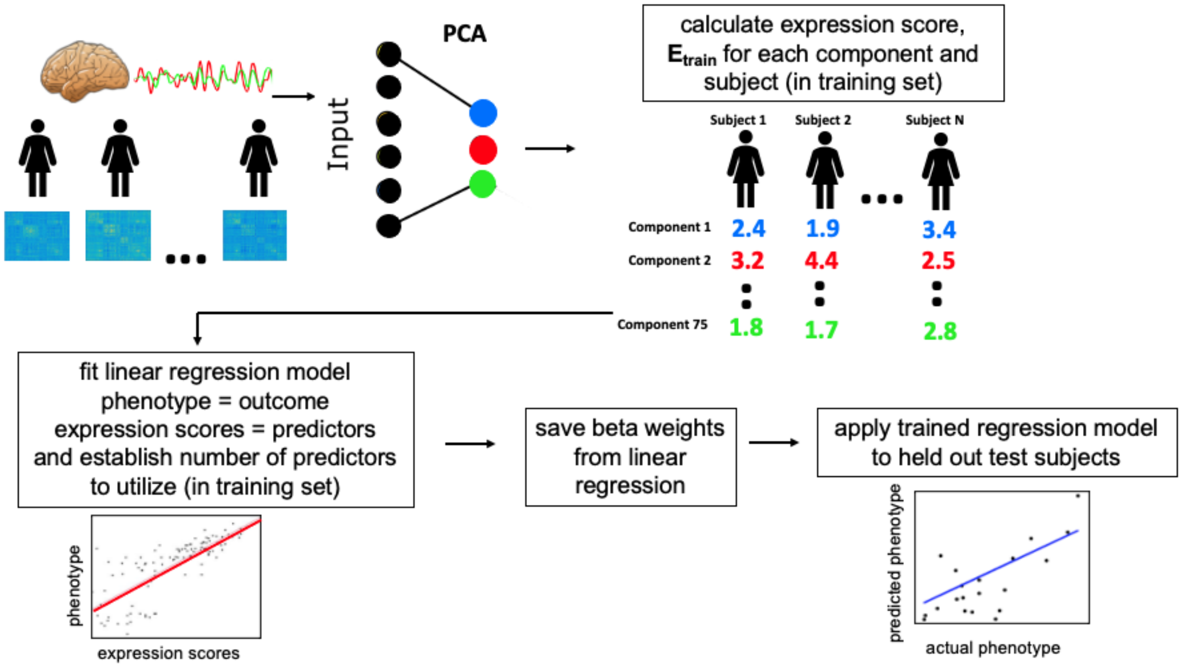


*Figure S2. Steps of Principal Component Regression Predictive Modeling.*

We implemented principal component regression (PCR) (7) as a multivariate predictive modeling method for identifying brain-behavior relationships (8) (see Figure S2). The method involves two key steps: 1) Use principal component analysis (PCA) to find a set of components that capture *inter-individual* differences in brain features; 2) Use multiple regression in a cross-validation framework to link expression scores for these components to phenotypes of interest. In previous work, we often used the more general name brain basis set (BBS) for this approach to capture commonalities with work by our group and others that use alternative methods for step 1 (e.g., independent component analysis (9,10) or community detection (11,12)). We chose the PCR approach for this study because our previous work showed it has high test-retest reliability (13) and predictive accuracy (14,15) and generally performs as well as or better than alternative methods such as support vector regression and ridge regression (13).

We performed PCA dimensionality reduction on an *n* subjects by *p* connectivity features matrix, yielding *n* principal components (i.e., directions in the feature space) that represent inter-individual differences in the imaging features (functional connectivity or graph theory metrics). Pre-subject expression scores for a subset of *k* of these components then entered multiple regression modeling to identify linear associations with phenotypes of interest (here, sleep duration). Of note, we selected *k* using 5-fold cross-validation within the training data, as in our previous work (13).

To assess accuracy and generalizability of PCR predictive models, we used leave-one-site-out cross-validation. In each fold of the cross-validation, data from one of the 21 sites served as the held-out test dataset and data from the other 20 sites served as the training dataset. Additionally, to ensure separation of train and test datasets, at each fold of the cross-validation, a new PCA was performed on the imaging features (functional connectivity or graph theory metrics) in the training dataset, and expression scores of these brain components were calculated for the test set. Note that by employing leave-one-site-out, members of twinships and sibships are never present in both training and test samples. We assessed the performance of PCR predictive models with cross-validated Pearson's correlation and cross-validated partial eta squared.

In each fold of the leave-one-site out cross-validation (LOSO-CV), PCR predictive models were trained in the train partition with the following covariates (unless explicitly stated otherwise for specific analyses): sex, race-ethnicity, age, age squared, mean FD and mean FD squared. To maintain strict separation between training and test datasets, regression coefficients for the covariates learned from the training sample were applied to the test sample to calculate effect size measures (Pearson's $\text{correlation}_{\text{cross-validated}}$). This procedure is described in detail in our previous publications (15,16).

We assessed the significance of all cross-validation-based correlations with non-parametric permutation tests. We randomly permuted the 5,596 participants' outcome variable values

10,000 times and reran the PCR predictive modeling stream at each iteration, yielding a null distribution of correlation values. The procedure of Freedman and Lane (17) was used to account for covariates. In addition, exchangeability blocks were used to account for twin, family, and site structure and were entered into Permutation Analysis of Linear Models (PALM) (18) to produce permutation orderings.

#### 6. Latent Variable Modeling for Sleep Duration

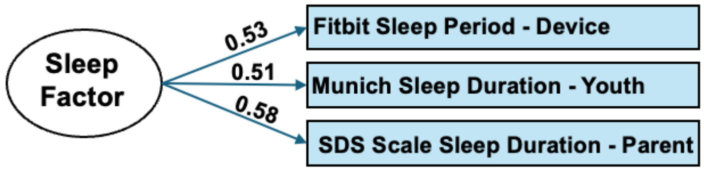


***Figure S3. Factor Model of Sleep Duration.*** *Path estimates reflect standardized factor loadings.*

To construct a latent variable for sleep duration that triangulates across objective (Fitbit) and subjective (parent- and youth-report) assessments, we conducted a factor analysis using the "fa" function from the psych package (version 2.4.6.26) in R (version 4.3.2). For the parent-reported Sleep Disturbance Scale (SDSC), we used the sleep duration item. This item asks "How many hours of sleep does your child get on most nights? 1 = 9-11 hours; 2 = 8-9 hours; 3 = 7-8 hours; 4 = 5-7 hours; 5 = Less than 5 hours. For each response, we assigned the midpoint of the associated range, and assigned 5 for the "Less than 5 hours response". For the youth-rated Munich Chronotype Questionnaire, we determined sleep duration as the difference between the mctq_sow_calc variable (Sleep onset, workday) and mctq_sd_wake_up_time_calc variable (Sleep end, workday). For the Fitbit data, we used the fit_ss_sleepperiod_minutes variable.

#### 7. Additional Socio-Environmental and Behavioral Variables

We examined associations between expression of a somatomotor disconnection graph theory component and a number of socio-environmental variables from the ABCD dataset. The number of participants included in this analysis varied depending on the availability of the variable. We provide additional information on these variables and the sample size for the analysis below.

*Household Income (n*=5,342) - This variable covered all sources of income for family members, including wages, benefits, child support payments, and others. It was assessed in bins as follows: 1 = <5,000, 2 = 5,000 - 11,999, 3 = 12,000 - 15,999, 4 = 16,000 - 24,999, 5 = 25,000 - 34,999, 6 = 35,000 - 49,999, 7 = 50,000 - 74,999, 8 = 75,000 - 99,999, 9 = 100,000 - 199,999, 10 = More than 200,000, and we assigned each subject the midpoint for their bin.

*Parental Education* (*n*=5,582) - This variable reflects the average educational achievement of parents or caregivers, and it is recoded into years based on the method of a previous ABCD report (19).

*Area deprivation index (n*=5,262) - This variable reflects an ABCD consortium-supplied variable (reshist_addr1_adi_wsum). Higher scores on the index indicate greater deprivation including higher percent of families living in poverty, increased unemployment, and lower levels of educational attainment at the neighborhood level.

*Child's screen time* (*n*=5,590) - Youth participants estimated the amount of time they spent on each of a series of six digital activities on both "typical" weekday and weekend days, which were then weighted (5*weekday + 2 * weekend) to yield a total screen estimate.

*Parent-reported externalizing and internalizing symptoms* (*n*=4,704) - Parents completed the *Child Behavior Checklist (CBCL)*. The CBCL includes two overarching scales. The Internalizing Problems composite scale is composed of the Withdrawn, Somatic Complaints, and Anxious/Depressed subscales. The Externalizing Problems composite scale is composed of the Delinquent Behavior and Aggressive Behavior subscales.

*Youth-reported externalizing and internalizing symptoms* (*n*=5,394) - Youth completed the *Brief Problem Monitoring Form*, an abbreviated 18-item version of the CBCL, which has internalizing and externalizing composite scores.

*Child's general cognitive ability (n*=4,445) - We created a general cognitive ability variable from bifactor modeling of the ABCD neurocognitive battery. Details on the creation of the variable and predictive validation of the variable are available in our previous studies in ABCD (16,20).

*Child's grades (n*=5,377) - Parents completed the *School Risk and Protective Factors Survey*, and child's grades were drawn from the *School Environment* subscale. We investigated youth-reported (sag_grades_last_yr) and parent-reported (sag_grade_type) overall grades over the last year. For each reporter, each response was assigned the midpoint of the associated range (e.g., a score of 1, indicating 97-100, was assigned the score of 98.5). We then calculated the average of the youth-reported and parent-reported overall grades as the final score.

##### 8. Comparison of the Somatomotor Disconnection Component Across Two Independent ABCD Samples

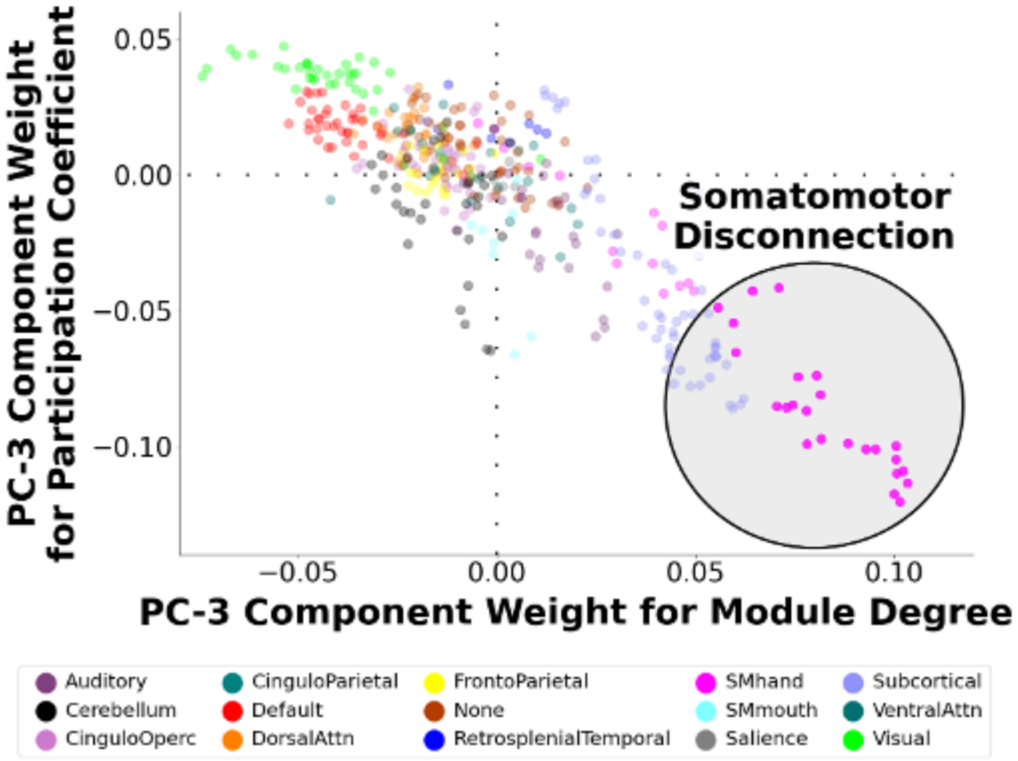


***Figure S4. Visualization of component #3 derived from the ABCD baseline sample. The component, which exhibits a prominent somatomotor disconnection motif, is highly similar to component #3 from the ABCD year 2 sample.***

We obtained component #3 from graph theoretic metrics from 3,148 participants from the ABCD baseline sample who have no overlap with the wave 2 participants used in the main analysis. We found that component #3 from this distinct sample was nearly identical to component #3 from the year-2 sample (see Figure S4), even though there was no overlap in the participants in the two samples. Indeed, feature weights for baseline- and year 2-derived components displayed a strong correlation of 0.99.
